# Colour pattern edge contrast statistics can predict detection speed and success at ecologically relevant viewing distances in triggerfish (*Rhinecanthus aculeatus*)

**DOI:** 10.1101/2022.06.16.496397

**Authors:** Cedric P. van den Berg, John A. Endler, Daniel E. J. Papinczak, Karen L. Cheney

## Abstract

Edge detection is important for object detection and recognition. However, we do not know whether edge statistics predict the detection of prey by non-human predators. Understanding the link between image statistics and animal behaviour is crucial and of increasing importance given the growing availability of image analyses and their application across non-human visual systems. Here, we investigated whether Boundary Strength Analysis (BSA), Local Edge Intensity Analysis (LEIA) and the Gabor Ratio (GabRat) could predict the speed and success with which triggerfish (*Rhinecanthus aculeatus*) detected patterned circular stimuli against a noisy visual background, in both chromatic and achromatic presentations. We found that individual pattern statistics could only explain up to 2% of the variation in detection time, whereas PCA regression analysis considering all edge statistics simultaneously explained up to 6% of the variation. This suggests that other factors explained more behavioural variation than individual edge statistics. Furthermore, different statistics significantly correlated with detection speed depending on treatment, viewing distance, and changes in fish response over time, while highlighting the importance of considering spatial acuity and relevant viewing distances in the study of visual signals. Our results demonstrate the need for broad and unbiased approaches for identifying task-specific predictive relationships between pattern statistics and animal behaviour using image statistics capturing different aspects of colour patterns. We require robust statistical approaches to investigate correlations between ecological effect and the ever-increasing dimensionality and size of datasets in the field of visual ecology, rather than pre-emptively narrowing down the choice of image statistics unless warranted by specific hypotheses.

**Summary statement:** Correlations between edge detecting colour pattern statistics and animal behaviour are complex. Specifically, correlations are unlikely to be explained by single image statistics and depend upon observer distance.

## 1. Introduction

Edge detection is crucial to the perception of spatial detail and informs cognitive processes such as object detection and discrimination (Bhagavatula et al., 2009; Cronin et al., 2014; Endler, 2006; Ruxton et al., 2018; Stevens and Cuthill, 2006). Edges, therefore, should have an important function in defensive animal colouration. For example, edges can allow animals to hide against visual backgrounds by disrupting an animal’s outline via disruptive camouflage (Cuthill et al., 2005; Endler, 2006; Troscianko et al., 2017). Alternatively, highly contrasting edges can help emphasize outlines of animals or body parts, helping to generate potent visual signals, such as those used for aposematic or deimatic signalling (Ruxton et al., 2018). Animals and objects with edge intensity distributions, frequencies, regularity and orientations matching those of the background tend to be difficult to detect or discriminate, whereas salient visual signals contrast against their visual background and are therefore easier to detect (Endler, 1978). In addition to informing object detectability *per se*, variation in edge contrast can have a profound impact on saliency and thus, search optimisation (Green et al., 2018; Krummenacher et al., 2010).

To approximate the perception of edge contrast at early stages of visual processing, colour pattern analyses relevant to animal vision can be performed using calibrated digital photography (Stevens et al., 2007), specifically using the Multispectral Image Calibration & Analysis (MICA) toolbox (Troscianko and Stevens, 2015) and its integrated frameworks such as Quantitative Colour Pattern Analysis (QCPA) (van den Berg et al., 2020b). There has been much work on quantifying various aspects of colour patterns including edge contrast, but there are few attempts to relate colour pattern statistics to ecologically relevant, task-specific behaviour using animal behaviour experiments. This problem is common in the study of defensive animal colouration, where the speed and reliability with which a predator can detect and locate prey is crucial in determining the survival rates of patterned prey, and hence the evolution of cryptic (Galloway et al., 2020) or conspicuous (Speed and Ruxton, 2005) defensive colouration.

In addition to the spectral sensitivities and relative abundance of photoreceptors of a visual system (Cronin et al., 2014), the perception of spatial detail and thus edge contrast depends on the acuity of an animal observer and the distance at which a visual signal is observed (Caves et al., 2016; Endler, 1978), which can dramatically alter the function of animal colouration. Despite the known species- and task-specificity of neuronal processing and cognition, few colour pattern statistics have been investigated for their ability to reflect ecological significance in a specific context for a particular animal observer. Investigations of how, and if, such modelled data correlates with animal behaviour are crucial, particularly given the steady increase in available image analyses and subsequent increase of data dimensionality.

Edge detecting colour pattern analyses in the QCPA include the Gabor Ratio Analysis (GabRat) (Troscianko et al., 2017), Boundary Strength Analysis (BSA) (Endler et al., 2018) and Local Edge Intensity Analysis (LEIA) (van den Berg et al., 2020b). Troscianko, Skelhorn and Stevens (2017) demonstrated that GabRat was more efficient in explaining detection speed of artificial grey scale moth stimuli in an achromatic search task for humans compared to 12 other edge detecting pattern metrics. However, GabRat has not yet been used in combination with spatial acuity and cone catch modelling assuming non-human observers. Sibeaux, Cole and Endler (2019) used the BSA to quantify female mate choice in Trinidad Guppies (*Poecillia reticulata)*. However, to our knowledge, there have been no studies investigating BSA in a predation context, specifically in relation to detection speed and success, nor in combination with spatial acuity and cone catch modelling. Lastly, while LEIA has been used in a study quantifying camouflage in precocial chicks (Rohr et al., 2021), no study, to our knowledge, has quantified correlations between LEIA statistics and animal behaviour.

To address these gaps, we investigated how QCPA edge detection analyses correlate with the response of a fish observer in a controlled experimental predation task. We investigated whether GabRat, BSA or LEIA could predict the speed and success with which triggerfish, *Rhinecanthus aculeatus* detect a circular stimulus of variable internal patterning against a noisy visual background. To do so we applied a range of analyses using both the investigation of individual statistics and dimensionality reduction analyses. We conducted two experiments with achromatic (treatment 1) and chromatic (treatment 2) stimuli to investigate possible differences in search performance due to additional noise from variation in colour in addition to luminance alone (Gegenfurtner and Kiper, 1992).

## 2. Material & Methods

### (a) Edge detecting pattern analyses in QCPA

BSA measures the colour and luminance contrast along edges, as well as the relative abundance of boundaries inside colour patterns (Endler et al., 2018). Similarly, LEIA quantifies colour and luminance edge contrast across an entire scene or object at the scale of individual edge detectors while providing a non-parametric approach to the measurement of edge distributions in an image (van den Berg et al., 2020b). GabRat was developed to reflect the functional principles of disruptive camouflage, quantifying the relative proportion and intensity of edges running orthogonally to the outline of an object (Troscianko et al., 2017). Here we investigated a total of 17 pattern edge statistics (BSA: 6; Gabrat, LEIA: 10; for a description of statistics used in this study see Table S1).

### (b) Study species

We used six adult triggerfish *Rhinecanthus aculeatus*, a common shallow reef inhabitant found throughout the Indo-Pacific, which feeds on algae, detritus and invertebrates (Randall et al., 1997). The species is easy to train and their visual system has been well-studied (Cheney et al., 2022). They have trichromatic vision based on a single cone (photoreceptor λ_max_ SW = 413 nm) and a double cone (photoreceptor λ_max_ MW = 480 nm and photoreceptor λ_max_ LW = 528 nm) (Cheney et al., 2013). The double cone members are used independently in colour vision (Pignatelli et al., 2010), but are thought to be used in tandem for luminance vision (Siebeck et al., 2014), as in other animals such as birds and lizards (Lythgoe, 1979). For this study, we have assumed both members to be responsible for luminance contrast perception (van den Berg et al., 2020a).

Fish were obtained from an aquarium supplier (Cairns Marine Pty Ltd, Cairns), shipped to The University of Queensland, Brisbane and housed in individual tanks of 120L (W: 40cm; L: 80cm, H: 40cm). Aquaria were divided in two halves by a removable black PVC partition. All animals had been housed at UQ for 2-4 years and used for previous behavioural experiments, which facilitated training with the animals having already learned to peck at visual stimuli for food. Experiments were conducted consecutively between September 2020 - February 2021.

### (c) Background design

Using a custom Matlab (MathWorks, 2000; version r2019b) script (originally written by JAE and modified by CvdB), a 14cm x 14cm noisy background was created on which target stimuli were displayed. The background was designed to mimic the spatial frequency distribution of natural marine habitats on a coral reef determined using images from Lizard Island (Great Barrier Reef), taken in February 2019 with a Nikon D810 in a Nauticam housing in depths of less than 3m, illuminated with natural sunlight. These images were then segmented using QCPA’s RNL clustering algorithm, using QCPA’s Gaussian acuity modelling to assume a triggerfish with an acuity of 3 cycles per degree (cpd) (Champ et al., 2014) observing the scenes at a distance of 10cm, a luminance JND threshold of 4 ΔS (van den Berg et al., 2020a) and a colour discrimination threshold of 2 ΔS (Green et al., 2022). The images were subjected to five cycles of RNL ranked filtering with a radius of 5 pixels and a falloff of 3. The resulting clustering was then used to determine the size distribution of randomly distributed clusters of 15,000 randomly shaped polygonal colour pattern elements belonging to 32 separate classes of equidistant 8-bit RGB values ranging from 0-0-0 RGB to 255-255-255 RGB for the achromatic and 0-0-0 RGB to 0-255-0 RGB for the chromatic background (Fig. 1).

**Figure 1:**
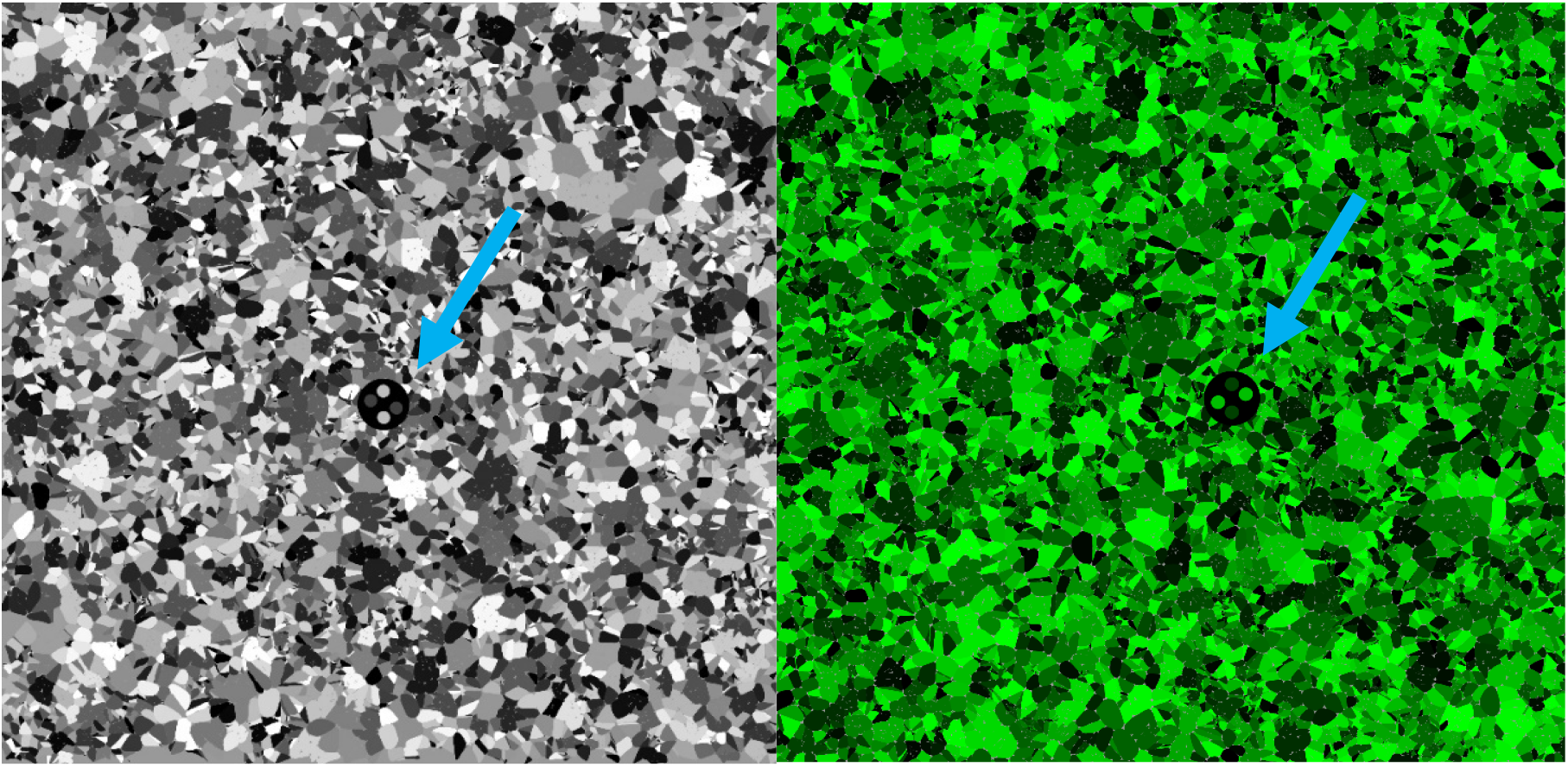
Examples of stimulus 1 randomly orientated on the noisy background for the achromatic (left) and chromatic (right) presentation. Blue arrows indicate stimulus position for illustrative purposes.

### (d) Stimulus design

We chose circles for the overall shape and symmetric circular internal patterning as these guaranteed equal numerical responses from the pattern analyses across trials, independent of rotation and viewing angle. Using repetitions of identical internal shapes further allowed for the numerical calibration of boundary contrast to theoretically be even across all stimuli when measured as the abundance weighted mean luminance contrast between pattern elements (BSA.BML, Table S1). This calibration allowed pattern contrast to vary within a mutual design constraint across all stimuli while allowing internal pattern variation. A set of four target stimuli and a training stimulus (diameter of 1cm) were developed using additional custom Matlab scripts (Table 1). Each stimulus comprised of a black background and a variable combination of internal patterning comprised of four regularly arranged smaller spots (diameter 0.25cm) (Table 1). Assuming a spatial acuity of 3 cycles per degree (cpd) (Champ et al., 2014), the internal spots would be discernible by the fish at distances below 4.8 cm, whereas the entire stimulus would be visible from as far as 19cm, with the distance from the divider separating fish from stimulus being 30cm.

**Table 1:**
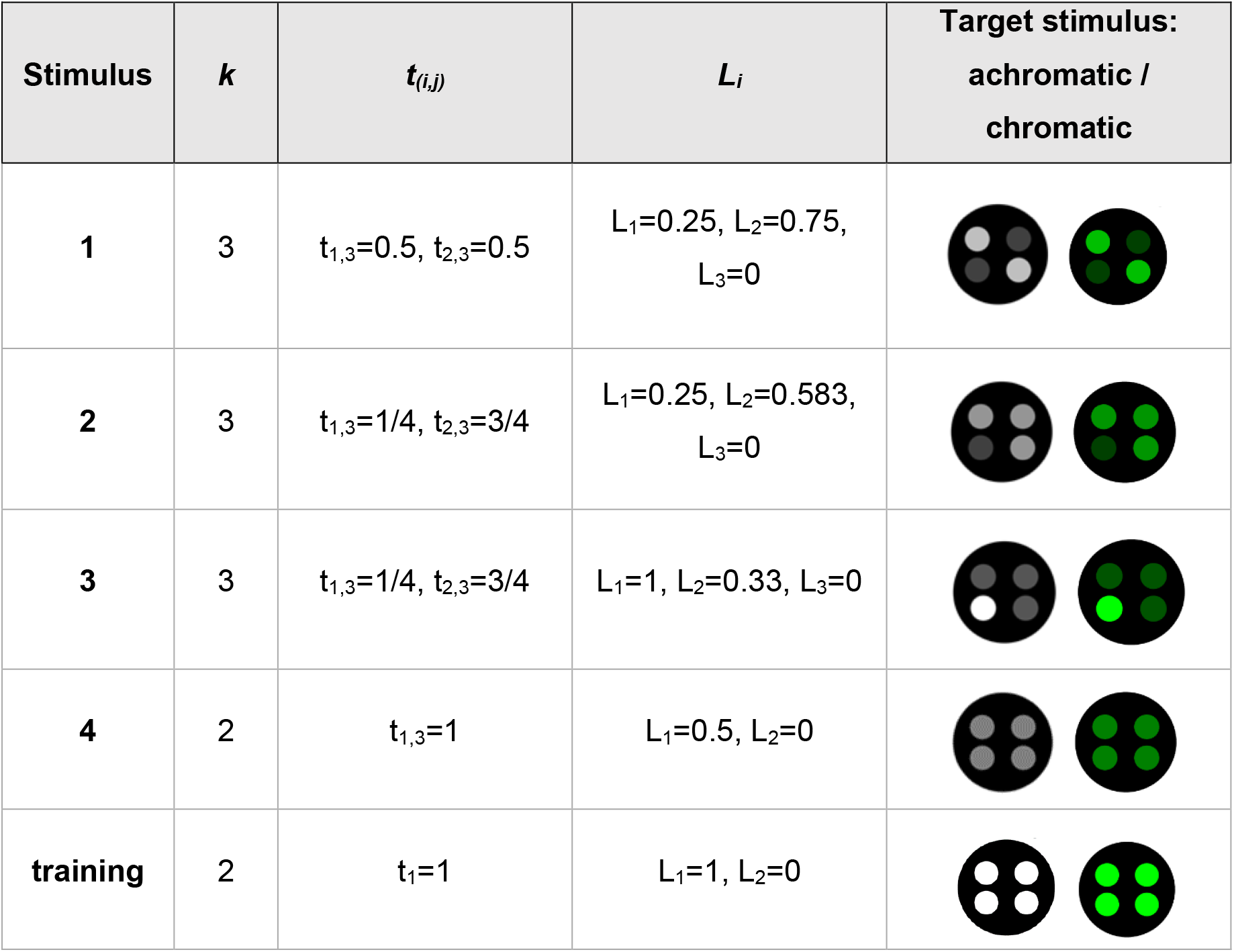
Parameters for stimuli used in the experiment. *K* = number of colour pattern elements, with element 3 being the black background in the spot. *t* = relative proportion of boundary type between elements *i* and *j*. L = luminance of each colour pattern element with L expressed as the % to max RGB (255) for the achromatic and % of G for the chromatic treatment.

| Stimulus | $k$ | $t_{(i,j)}$ | $L_i$ | Target stimulus:<br>achromatic /<br>chromatic |
| --- | --- | --- | --- | --- |
| 1 | 3 | $t_{1,3}=0.5, t_{2,3}=0.5$ | $L_1=0.25, L_2=0.75,$<br>$L_3=0$ | |
| 2 | 3 | $t_{1,3}=1/4, t_{2,3}=3/4$ | $L_1=0.25, L_2=0.583,$<br>$L_3=0$ | |
| 3 | 3 | $t_{1,3}=1/4, t_{2,3}=3/4$ | $L_1=1, L_2=0.33, L_3=0$ | |
| 4 | 2 | $t_{1,3}=1$ | $L_1=0.5, L_2=0$ | |
| training | 2 | $t_1=1$ | $L_1=1, L_2=0$ | |

For the achromatic treatment, the internal dots were grey, with equal RGB stimulus values for each pattern element (e.g. 80-80-80 RGB). For the chromatic (green) treatment, the same patterning was repeated, but the R and B pixel value remained fixed at 0 while the G value was identical to the achromatic treatment.

### (e) Stimulus placement and session design

Using a third custom Matlab script, each stimulus was placed on the respective background at nine positions (top-left, top-middle, top-right, middle-left, middle-middle, middle-right, bottom-left, bottom-middle, bottom-right) in a random orientation (Fig. 1). We then compiled sessions of six stimulus presentations, using five pseudo-randomly drawn stimuli and position combinations from all possible stimulus and position combinations. To balance each session, the sixth stimulus and position combination for each session was determined by pseudo-randomly choosing from the list of least represented positions and stimuli of a given session. This session design was repeated to achieve a minimum of 30 repetitions of each stimulus for each animal (mean = 38.75, SD = 4.04) consisting of at least 3 repetitions of each position for each stimulus (mean = 4.16, SD = 1.30) across all achromatic sessions (n = 27) and chromatic sessions (n = 29). To ensure each stimulus ended up being tested equally as often for all animals and positions by the end of trials, stimulus and position frequencies were tested using a Chi-square test. Stimuli were displayed on an iPad Air 2 fitted with a matte screen protector and placed in a waterproof case (Lifeproof Nuud iPad case) with brightness set to maximum. The iPad was then placed parallel to the aquarium floor 10cm from the bottom and against the back wall.

### (f) Stimulus quantification

Image analysis was performed with the MICA toolbox (version 2.2) running on ImageJ (version 1.53a) using a custom designed automated batch script of the QCPA. To quantify the edge contrast provided by each of the three pattern statistics (BSA, GabRat, LEIA), a calibrated Olympus E-PL5 Penlight camera with a 60mm macro lens was used to take images of each of the nine replicated positions for each stimulus and treatment. The pictures were taken in the dark (as LED screens emit rather than reflect light) and the brightest patch class (white or bright green) used to calibrate each image using the ‘estimate black point’ function when creating the normalised and standardised multispectral image files (.mspec). The ‘white’ patches (255-255-255 RGB) were determined to be of 72.5% reflectance (even though technically radiance, but the input to QCPA is in reflectance), comparing the radiance of the patches to the reflectance of a Spectralon (Ocean Optics) white standard illuminated by a PX-2 light source (Ocean Optics). A chromatic cut-off (average cone catch per pixel below which no chromaticity is possible) was set at 3%, preventing artificial chromaticity due to minor absolute differences between cones.

Each image (n = 36) was manually segmented into stimulus and corresponding background using manual image segmentation in ImageJ (Schneider et al., 2012) and colour patterns analysed using an automated QCPA script. Each image was analysed at a modelled viewing distance of 2cm, 5cm, 10cm and 30cm using the Gaussian acuity modelling function in QCPA, resizing the images to a pixel per minimally resolvable angle (MRA) ratio of 5, thus removing spatial detail that cannot be resolved at a given distance (van den Berg et al., 2020b). The viewing distances are within the range of distances encountered by the fish from pecking a stimulus to observing the stimulus from the divider (30cm). GabRat analysis was performed assuming a 1 cpd peak contrast sensitivity function (CSF) acuity based on the CSF curve shape of a Black-faced blenny (*Triperygion delaisi*) (Santon et al., 2019), the only marine fish for which a CSF is available. Despite the distant relationship, the general shape of the CSF is representative of most vertebrates, peaking at around a third of the maximum (da Silva Souza et al., 2011). For LEIA, the images were further subjected to a 5 times RNL-ranked filtering with a radius of 5 pixels and a falloff value of 3 to remove artificial colour gradients introduced during the acuity modelling. LEIA values were obtained from the untransformed edge histograms. For BSA, the images were further subjected to RNL clustering (Fig. S1), using a 2 JND chromatic and 4 JND achromatic threshold which were based on empirical findings in past studies (Green et al., 2022; van den Berg et al., 2020a). RNL contrast was determined by using Weber fractions of 0.07:0.05:0.05:0.05 for sw:mw:lw:dbl, spectral sensitivities and a white LED illuminant spectrum as per (van den Berg et al., 2020a). Weber fractions were calculated assuming a receptor noise of 0.05 and a relative cone abundance of 1:2:2:2 (sw:mw:lw:dbl).

### (g) Animal training

Using operant conditioning, fish were trained to peck at a piece of squid placed on a black spot (diam. 1cm) randomly placed (using natural adhesive properties) on a uniform grey background displayed on an iPad. Once fish had pecked at the food on the target, they were given a second piece of squid from above with tweezers. The size of the food reward on the target spot was subsequently reduced until the fish were pecking at the target spot without any food on it. Next, the target was changed to a patterned spot (Table 1, stimulus 1) on a plain background and, finally, on a noisy background (Fig. 1). Fish moved to each training stage when successful in >80% trials over 6 consecutive sessions of 6 trials per day. A trial was considered unsuccessful if the fish took longer than 30 seconds (measured using a stopwatch) after swimming through the door of the divider to make a choice or if it pecked at the background more than twice. As the fish sometimes get distracted, we allowed fish two wrong pecks before concluding that fish had not detected the target. Testing was suspended for the day if the fish showed multiple timeouts (failure to peck at the stimulus within 30 seconds). Once fish had completed treatment 1 (achromatic stimulus), they were re-trained for treatment 2 (chromatic treatment) and had to meet training criteria prior to commencing trials.

### (h) Animal testing

For both treatments, six individuals were tested. However, two individuals did not complete the chromatic trials and were excluded from the chromatic data analysis. Every fish was tested for one session of six trials per day, with each session determined as described above. Stimulus 1 did not get drawn for the first three sessions in the achromatic treatments (Fig. 2). As per training, a trial was considered unsuccessful if the fish took longer than 30 seconds to make a choice or if it pecked at the background more than twice. Time to detection was recorded as the time between the moment the fish moved past the divider and the successful peck at the target spot.

**Figure 2:**
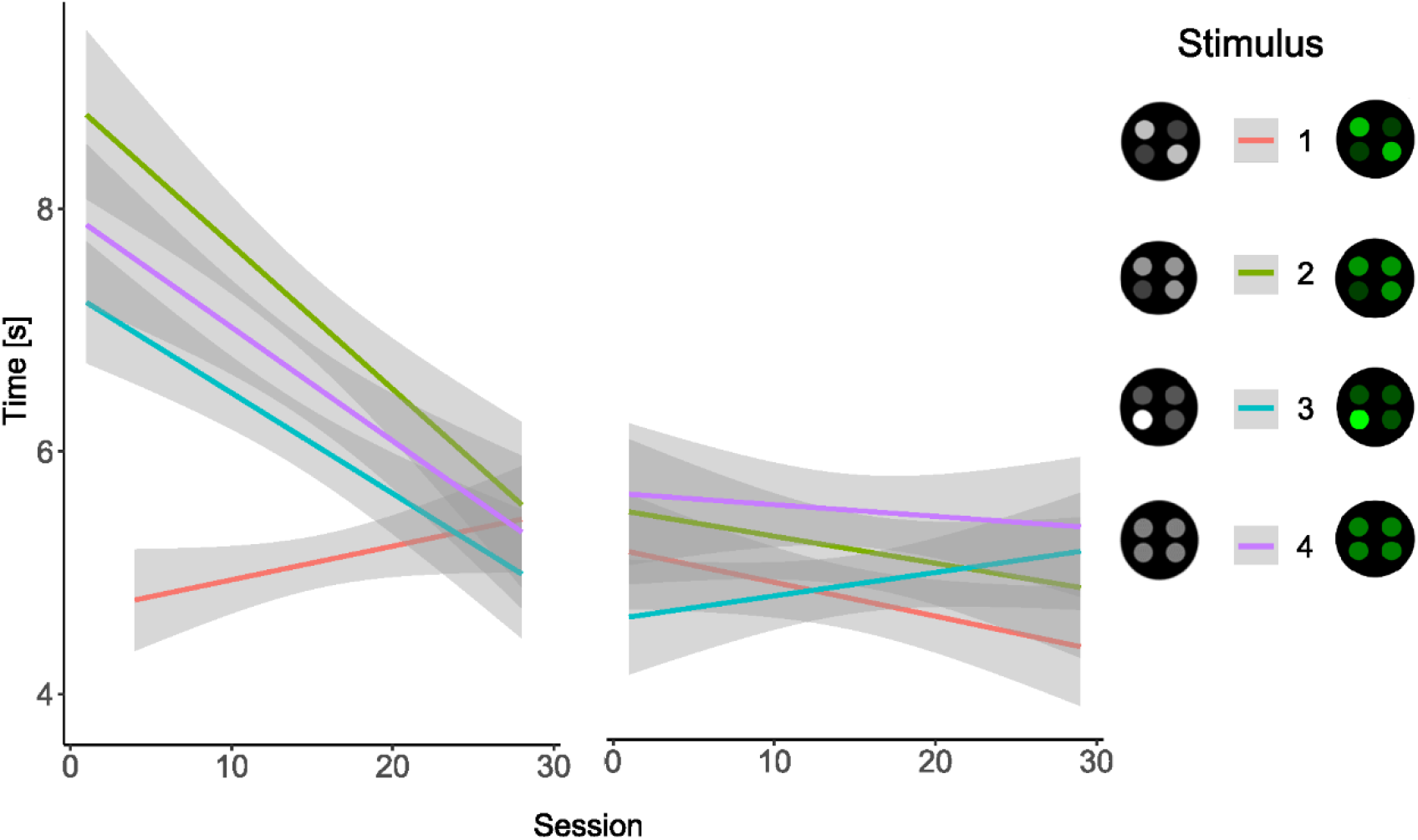
Marginal effects plot of the detection time [s] for each session and stimulus summarised across all animals (achromatic (left): n=6, chromatic (right): n=4), 95% confidence intervals indicated with shadows.

### (i) Statistical analysis

All statistics were performed in R (R Core Team, 2021; V4.0). Time to detection was left skewed and was transformed using an ordered quantile (QRQ) normalizing transformation with the *bestNormalize* package (Peterson and Cavanaugh, 2020) prior to subsequent analyses to ensure normality. Pattern statistics were normalised using the preProcess function in the *caret* package (Kuhn, 2021; v6.0-88); with the “range” option to restrict values to range from 0 to 1. To ensure the image statistics at each viewing distance (2cm, 5cm, 10cm, 30cm) were capturing the pattern differences between the stimuli, we used regularized discriminant analysis (RDA) (Friedman, 1989) to predict the stimulus category. RDA classification was done using the *caret* and *klaR* (Weihs et al., 2011) R packages. This confirmed that the selected pattern statistics were effectively delineating the stimuli at all viewing distances with the stimulus type prediction rates of the RDA trained model being 100% at all viewing distances.

Differences in time to detection between stimuli of each treatment were tested with a linear mixed effects model accounting for session number as a fixed effect (after confirming the absence of an interaction with stimulus) and fish ID as a random effect using the lmer function on the lme4 package (Bates et al., 2015). Failure rates between stimuli were compared with a Fisher-exact test in the *rstatix* package (Kassambara, 2021; v0.7.0).

We then investigated whether colour pattern metrics correlated with detection speed. For BSA only the original BSA statistics provided by the QCPA was used to capture luminance and chromatic contrast (Table S1). For GabRat, only the achromatic (dbl cone) contrast was used, as the acuity for luminance contrast detection was assumed to be dominating the acuity provided by the chromatic channels (Lind and Kelber, 2011). We looked at both, the statistics of the stimuli by themselves as well as in contrast to their visual backgrounds. This was done by using the absolute difference between a given stimulus statistic and the corresponding background. As GabRat measures the appearance of a stimulus edge against its background, GabRat values were identical in both instances.

To identify each pattern statistic’s ability to predict detection speed, the QRQ transformed time to detection for each pattern statistic was investigated by fitting a linear mixed effect model using the lmer function in the lme4 package. Fish ID was added as a random effect to account for individual differences with viewing distance as a nested random effect. The amount of deviance in the time to detection explained by each pattern statistic was quantified using the pamer function in the *LMERConvenience Functions* R package (Tremblay, 2020; v3.0). We omitted applying alpha corrections as per Troscianko, Skelhorn and Stevens (2017) to prevent inflating type II errors. The position of the stimulus and the trial number were included as fixed effects with an interaction term to reduce the amount of unexplained variation in each model and to account for learning effects.

A principal component regression analysis (PCR) was applied to all pattern statistics at each viewing distance. The PCR was done using the *pls* package (Bjorn-Helge et al., 2020) to find a set of principal components which explains a maximum amount of variation in detection speed while maximising parsimony. To identify the best combination of fully weighted predictors, a stepwise regression analysis with sequential replacement was conducted using the *leaps* package (Lumley and Miller, 2020) to identify the model with the lowest prediction error. PCA regression analysis and RDA were performed by randomly selecting 80% of the data as training data and 20% as test data.

## 3. Results

### (a) Differences in detection speed and success between stimuli

#### Treatment 1 (achromatic stimuli)

We conducted a total of 866 achromatic trials, of which fish successfully detected the target stimuli in 809 trials. There were no individual differences in performance across stimuli regarding the failure rate (*p* = 0.955); however, failure rates between stimuli varied significantly (Stim 1: 3.58% or 8 out of 223, Stim 2: 9.76% or 20 out of 205, Stim 3: 3.18% or 7 out of 220, Stim 4: 10.10% or 22 out of 218; p = 0.004).

Detection speeds varied significantly between stimuli (F_796.21_ = 4.69, *p* = 0.003) with stimulus 1 being detected significantly faster than stimulus 2, 3 and 4. Detection times improved over time for stimulus 2 (F_179.53_ = 9.55, *p* = 0.002), stimulus 3 (F_206.41_= 7.39, *p* = 0.007) and stimulus 4 (F_194_ = 10.87, *p* < 0.001), but not for stimulus 1 (F_208.49_ = 1.75, *p* = 0.10) (Fig. 2). A total of 57 out of 866 stimuli failed to be detected.

#### Treatment 2 (chromatic stimuli)

We conducted a total 692 chromatic trials, of which fish successfully detected the target stimuli in 677 trials. Fail rates in the chromatic treatment were lower than fail rates in the achromatic treatment (2.17% chromatic failure rate vs. 6.58% achromatic fail rate measured as failed trial proportion of all trials). There were no individual differences in performance across stimuli regarding the failure rate (*p* = 0.212). While failure rates between stimuli did not vary significantly (*p* = 0.176)) the pattern of fail rates between stimuli resembles the achromatic treatment with stimuli 1 & 3 having the lowest fail rates (Stim 1: 0.58% or 1 out of 172, Stim 2: 4.10% or 7 out of 171, Stim 3: 1.69% or 3 out of 177, Stim 4: 2.33% or 4 out of 172).

Detection times for the chromatic context did not vary between stimuli (F_666.06_ = 2.01, *p* = 0.11). However, while detection times generally improved over time (F_669_ = 4.56, *p* = 0.03), individual stimuli did not: Stimulus 1 (F_166.03_ = 0.84, p = 0.36), Stimulus 2 (F_159.75_ = 0.86, p = 0.36), Stimulus 3 (F_169.16_ = 0.32), Stimulus 4 (F_163.51_ = 2.47, p = 0.12).

### (b) Investigating individual pattern statistics to predict detection speed

The amount of explained variation in detection speed varied substantially between analyses, but overall was very low. Significant single-statistic analyses considering viewing distance as a random factor were rarely able to explain more than 1% of variation (max: 1.22%, min: 0.06%, mean: 0.35%, std: 0.29, Table 2). Considering the viewing distances separately yielded larger proportions of explained variation (max: 1.65%, min: 0.24%, mean: 0.82%, std: 0.37, Fig. 3 & 4, see table S2 for details).

**Table 2:** Summary table of the achromatic, chromatic and combined results of the proportion of deviance explained by each model tested for each image statistic with viewing distance as a nested random effect. Only statistics with significant interactions are listed (out of 17, see Table S1).

| Stimulus only (combined) |  |  |  | Animal vs. Background (combined) |  |  |  | Stimulus only (chromatic) |  |  |  |
| --- | --- | --- | --- | --- | --- | --- | --- | --- | --- | --- | --- |
| Pattern statistic | F-value | p-value | % expl. dev. | Pattern statistic | F-value | p-value | % expl. dev. | Pattern statistic | F-value | p-value | % expl. dev. |
| BSA.BCVL | 53.45 | <0.001 | 0.80 | BSA.BCVL | 67.31 | <0.001 | 1.00 | GabRat | 16.19 | <0.001 | 0.53 |
| Col.CoV | 38.57 | <0.001 | 0.57 | BSA.BML | 33.65 | <0.001 | 0.50 | Lum.kurt | 15.79 | <0.001 | 0.51 |
| Col.mean | 67.80 | <0.001 | 1.00 | Col.CoV | 36.71 | <0.001 | 0.55 | Col.sd | 10.65 | 0.001 | 0.34 |
| Col.sd | 80.80 | <0.001 | 1.19 | Col.mean | 78.36 | <0.001 | 1.15 | Lum.skew | 9.55 | 0.002 | 0.31 |
| Lum.CoV | 45.49 | <0.001 | 0.68 | Col.sd | 82.92 | <0.001 | 1.22 | Col.skew | 7.98 | 0.005 | 0.26 |
| Lum.kurt | 13.43 | <0.001 | 0.20 | Lum.CoV | 34.72 | <0.001 | 0.52 | Lum.CoV | 6.96 | 0.008 | 0.23 |
| BSA.BML | 11.09 | 0.001 | 0.17 | Lum.kurt | 23.42 | <0.001 | 0.35 | Col.kurt | 6.73 | 0.010 | 0.22 |
| Col.kurt | 10.80 | 0.001 | 0.16 | Lum.skew | 24.53 | <0.001 | 0.37 | Col.CoV | 4.96 | 0.026 | 0.16 |
| Lum.skew | 8.12 | 0.004 | 0.12 | BSA.BsSsat | 15.50 | <0.001 | 0.23 |  |  |  |  |
| BSA.BsSsat | 7.65 | 0.006 | 0.11 | BSA.BsL | 12.38 | <0.001 | 0.19 |  |  |  |  |
| Col.skew | 6.41 | 0.011 | 0.09 | GabRat | 5.81 | 0.016 | 0.09 |  |  |  |  |
| Lum.mean | 6.32 | 0.012 | 0.09 | BSA.BCVSsat | 4.88 | 0.027 | 0.07 |  |  |  |  |
| GabRat | 5.81 | 0.016 | 0.09 | Lum.mean | 4.11 | 0.043 | 0.06 |  |  |  |  |
| BSA.BsL | 5.38 | 0.020 | 0.08 |  |  |  |  |  |  |  |  |
| Stimulus only (achromatic) |  |  |  | Animal vs. Background (achromatic) |  |  |  | Animal vs. Background (chromatic) |  |  |  |
| Pattern statistic | F-value | p-value | % expl. dev. | Pattern statistic | F-value | p-value | % expl. dev. | Pattern statistic | F-value | p-value | % expl. dev. |
| BSA.BCVL | 22.09 | <0.001 | 0.61 | Col.mean | 27.98 | <0.001 | 0.76 | Col.sd | 15.36 | <0.001 | 0.50 |
| Col.mean | 26.83 | <0.001 | 0.73 | Col.sd | 22.73 | <0.001 | 0.62 | GabRat | 16.19 | <0.001 | 0.53 |
| Col.sd | 30.46 | <0.001 | 0.83 | Col.kurt | 15.89 | <0.001 | 0.43 | Lum.skew | 8.29 | 0.004 | 0.27 |
| Lum.CoV | 19.82 | <0.001 | 0.55 | BSA.BCVL | 9.02 | 0.003 | 0.25 | Lum.kurt | 7.45 | 0.006 | 0.24 |
| Col.kurt | 16.04 | <0.001 | 0.44 | Col.skew | 8.26 | 0.004 | 0.23 | Col.mean | 6.33 | 0.012 | 0.21 |
| BSA.BCVSsat | 10.32 | 0.001 | 0.29 | Lum.CoV | 7.65 | 0.006 | 0.21 | Col.skew | 5.64 | 0.018 | 0.18 |
| Col.skew | 9.52 | 0.002 | 0.26 | GabRat | 5.58 | 0.018 | 0.15 | Lum.CoV | 5.53 | 0.019 | 0.18 |
| BSA.BMSsat | 9.15 | 0.003 | 0.25 | BSA.BsL | 4.47 | 0.035 | 0.12 | Col.CoV | 5.35 | 0.021 | 0.18 |
| BSA.BsL | 7.84 | 0.005 | 0.22 | Lum.skew | 4.34 | 0.037 | 0.12 | BSA.BCVL | 4.68 | 0.031 | 0.15 |
| Lum.sd | 6.66 | 0.010 | 0.18 | Lum.kurt | 4.24 | 0.040 | 0.12 | BSA.BsL | 4.51 | 0.034 | 0.15 |
| GabRat | 5.58 | 0.018 | 0.15 | Lum.sd | 3.88 | 0.049 | 0.11 | BSA.BsSsat | 4.46 | 0.035 | 0.15 |
| Lum.kurt | 4.47 | 0.035 | 0.12 |  |  |  |  | Col.kurt | 4.41 | 0.036 | 0.14 |

**Figure 3:**
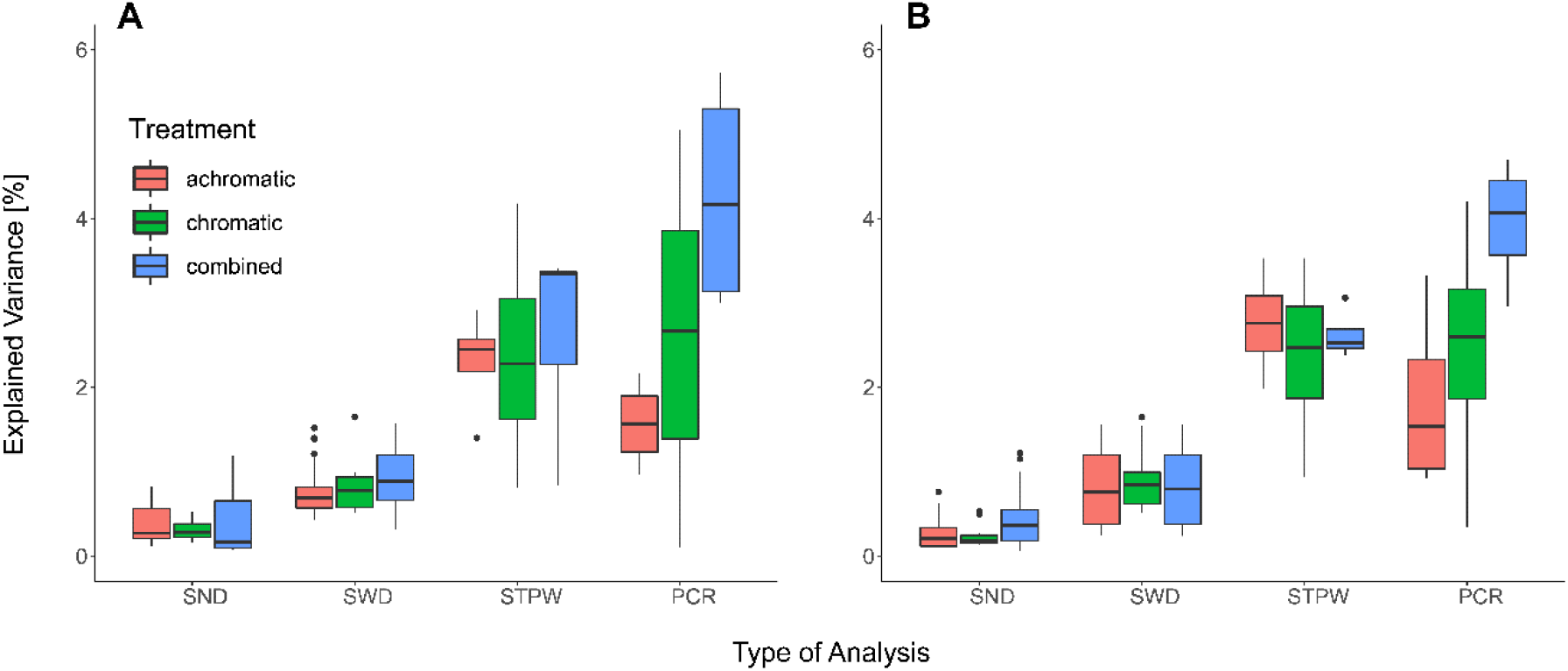
A) Explained variation in detection speed explained by the different analyses using only the stimulus pattern statistics, and B) using the difference between the stimulus and its respective background. SND: All significant single statistics with distance as random effect, SWD: all significant single statistics for each viewing distance, STPW: Stepwise regression analysis for each viewing distance, PCR: Principal component regression analysis for each viewing distance.

Considering all possible pattern statistics provided more explained variation with the stepwise regression analysis able to explain 2-4% (max: 4.18%, min: 0.81%, mean: 2.52%, std: 0.85, Fig. 3) whereas the principal component regression analysis was able to explain up to 6% (max: 6.33%, min: 0.11%, mean: 2.93%, std: 1.69, Fig. 3), with statistics of all three pattern analyses represented in all analyses.

However, different sets of pattern statistics were relevant at different viewing distances (Fig. 4, see tables S2-S4 for details). There was seemingly little difference between the use of the stimulus statistics themselves and the use of the respective stimulus-background contrast considering the average explained variation in fish behaviour. However, there are substantial differences between considering the stimulus by itself or in the context of its background when considering each statistic individually (Fig. 4), with maximum explained variability in detection speed coinciding with the estimated limit of internal stimulus pattern perception (∼5cm) for most comparisons (but see Fig. 4 D).

**Figure 4:**
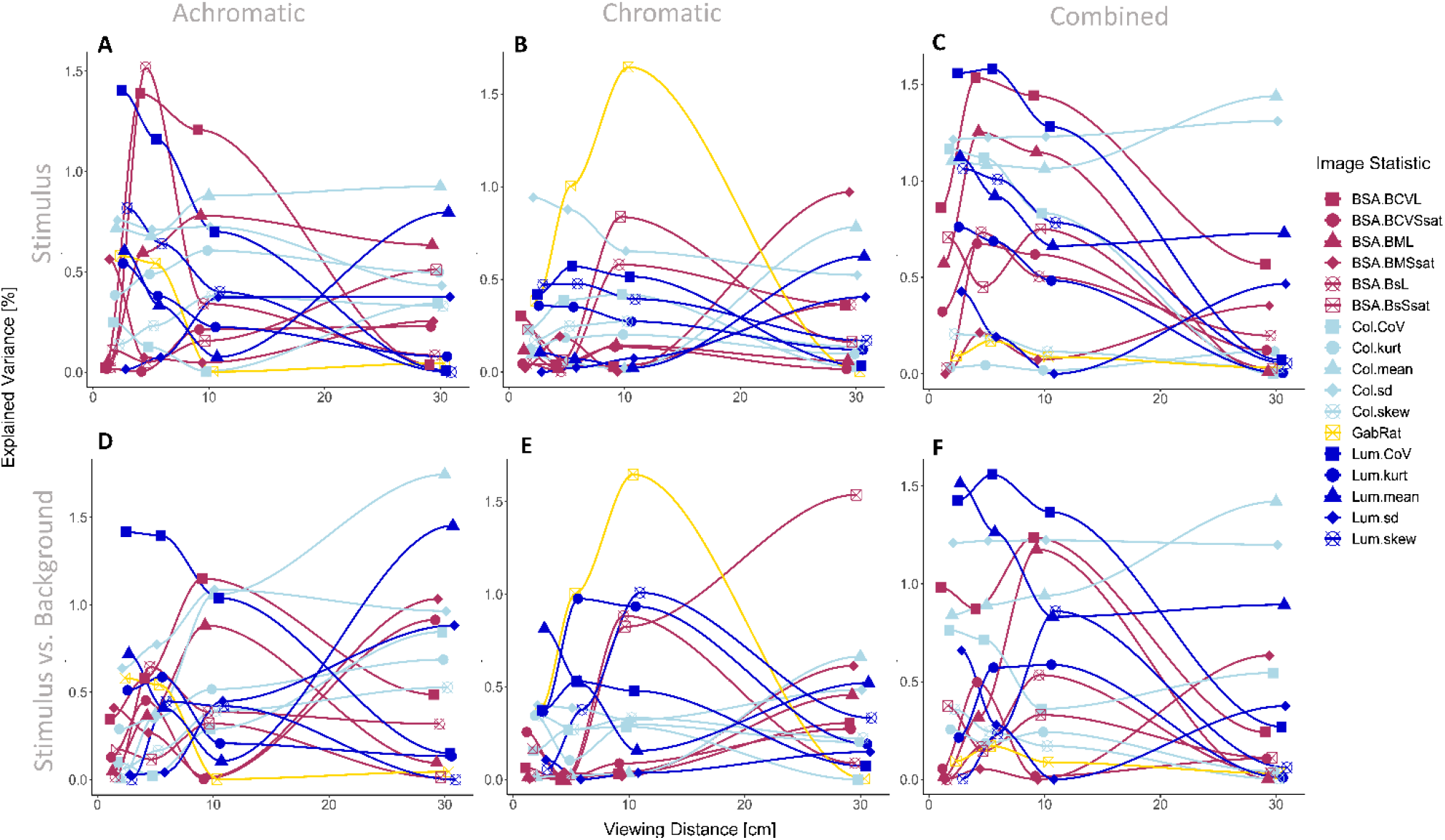
Explained variation in detection speed by each statistic at each distance. A) achromatic treatment, considering only stimulus statistics; B) chromatic treatment, considering only stimulus statistics; C) combined data of both treatments, considering only stimulus statistics; D) achromatic treatment, considering stimulus against background statistics; E) chromatic treatment, considering stimulus against background statistics; F) combined data of both treatments, considering stimulus against background statistics.

## 4. Discussion

We show that all edge detecting colour pattern analyses in the QCPA framework (BSA, LEIA, GabRat) can contribute to the prediction of stimulus detectability. However, the explained variation in detection speeds was low, but similar to that of most statistics investigated by Troscianko, Skelhorn and Stevens (2017) for human viewers. This confirms that stimulus detectability likely is subject to a complex array of factors, with edge contrast being one of many. Rather than focusing on a select few image statistics from the start, our results suggest that starting off with an unbiased presumption on the potential validity of a large array of pattern statistics can be a valid and objective approach towards identifying key morphological features contributing to the behavioural outcome under investigation. We further show that differences in internal patterning of visual stimuli lead to significant changes in search optimisation, detection speed and detection success. While our study focuses on a detection task in the context of a specific set of edge detecting pattern statistics, it is reasonable to assume that different types of visual information and cognitive processes are relevant for observed or assumed behavioural and ecological outcomes. These properties are reflected by different, task-specific, constellations of pattern statistics of which this study only considers a few.

We show that some aspects of the visual appearance are important in determining detection speed at certain distances, while not being relevant at others. This is intuitive, as brightness, colour, and pattern geometry change differently as a function of viewing distance, spectral sensitivities, photoreceptor abundances and discrimination thresholds. Our results align with previous results that estimating a few statistics of early-stage visual processing only reflects a fraction of the visual and cognitive processing underlying the ultimate behaviour (Troscianko et al., 2017). As viewing distances increase, finer internal patterns start to blur, subsequently changing the appearance of both the stimulus and its internal patterning, as well as the visual background. As a result, various mechanics captured by the pattern analyses used in this study, change, leading to a variable landscape of distance dependent correlations between pattern statistics and animal behaviour (Fig. 4). However, feature detectors in visual systems operate at different spatial scales (Elder and Sachs, 2004; Hubel and Wiesel, 1962). Such variable feature detection at a given distance could contribute to our understanding of differences in acuity estimates derived from behavioural experiments as opposed to histology; or between behavioural experiments using different sets of stimuli and paradigms and different viewing distances.

We further show that the average correlation of pattern statistics with detection speed is similar when considering the stimuli by themselves or with their respective visual backgrounds. However, there are large relative differences in behaviour prediction success between pattern statistics and viewing distances depending upon whether one considers the background or not (Fig. 4). Indeed, tactical signal design is intrinsically linked to the perception of a signal in the context of its visual background (Guilford & Dawkins, 1991). Therefore, it is inappropriate to ignore the backgrounds in studies where visual signals are observed against a multitude of different visual backgrounds (Niskanen and Mappes, 2005; Pike, 2018). We expect stronger differences between prediction success with and without backgrounds when conducting similar experiments with highly variable visual backgrounds.

Our data show a pronounced effect of increasing detection speed over time in the achromatic treatment but not in the chromatic treatment. However, the effect is not equally as strong for all stimuli. While stimuli 2, 3 and 4 significantly improved detection speeds over time, stimulus 1 was detected at maximal speed from the beginning (Fig. 2), indicating adaptation in how efficiently the animals were able to detect certain stimuli. This could be explained by a gradual change in search pattern (Credidio et al., 2012) coinciding with shifts in selective attention to specific features (Langley et al., 1996), a crucial effect of colour pattern diversity on predator cognition and a key mechanism behind apostatic selection as well as the evolution and persistence of colour pattern variability and polymorphism in nature (Bond and Kamil, 2006).

Differences in detection speed could also be due to the presence of a distinct visual feature unique to stimulus 1, making it significantly easier to detect. While stimulus 1 did not get presented in the first three trials, the response is profoundly different to that of the other stimuli despite a similar difference in internal contrasts and, importantly, persists throughout the duration on the achromatic trials (Fig. 2). This makes the difference in animal response unlikely to be the consequence of a novelty effect due to delay in presentation. While the range of contrast intensity in stimulus 1 is not unique, the diagonal symmetry and potential presence of a diagonal symmetry axis is (Fig. 1). The salience of this axis could be explained by mechanisms of perceptual grouping (Brooks, 2014) and/or the presence of feature detectors with differential selectivity to stimulus orientation (Hubel and Wiesel, 1962). Perceptual grouping is crucial to the strategic design of defensive colouration, such as the function of disruptive colouration (Espinosa and Cuthill, 2014) as well as background matching (Dimitrova and Merilaita, 2012). Unsurprisingly then, perceptual grouping can also aid in the detection of patterned prey, adding emphasis to the potential importance of symmetry in salient visual signals, especially when seen against an irregular background (Forsman and Merilaita, 1999; Forsman and Merilaita, 2003). However, we are not aware of any existing colour pattern analyses capable of quantifying ‘illusory’ features created by perceptual grouping in both, human and non-human observers. Therefore, this remains an intriguing area of investigation for future research.

The absence of improved detection over time does not explain the reduced error rate for stimulus 3 which is equally as low as for stimulus 1. Interestingly, stimulus 3 features a single high contrast marking, distinguishing it from stimuli 2 and 4 (Table 1). Thus, while not making the stimulus easier to detect (i.e. by aiding in switching from sequential to parallel search (Sagi and Julesz, 1984)), the vivid marking potentially helps identifying the stimulus upon detection. This may highlight cognitive differences between object detection and recognition and thus tactical and strategic signal design (Guilford and Dawkins, 1991; Hebets and Papaj, 2005). This is relevant in studies investigating the ecological significance of animal colouration, as well as studies investigating psychophysical thresholds (e.g. Santiago *et al*., 2020; van den Berg, Hollenkamp, *et al*., 2020). Given the salient markings in Stimulus 1, the coinciding reduction in error rates across both treatments makes sense although we cannot delineate whether this is caused by facilitated detection, discrimination, or both.

Developing approaches to the analysis of high-dimensional visual modelling data is a key requirement to the investigation of colour pattern space (van den Berg et al., 2020b). By using a variety of dimensionality reduction approaches, we highlight a subset of tools which can be used to navigate high-dimensional datasets such as those provided by the QCPA. The processing of visual signals from the moment information is registered by photoreceptors to the moment a behavioural response is observed is variable in specificity and complexity. Consequently, this makes it reasonable to assume anything from a single pattern statistic to complex multiparameter interactions to correlate with ecologically relevant animal behaviour.

Our study merely investigates 17 out of > 200 QCPA image statistics, all of which exclusively measure edge contrast. This array of statistics captures only a very limited set of visual features. Different pattern statistics capture different aspects of visual signals which are affected differently by natural selection. Consequently, high levels of correlation between specific colour pattern statistics and animal behaviour can be found in one context and not another. For example, while highly significantly correlated, GabRat by itself managed to barely explain 1% of variation in animal behaviour in this specific experiment (Table 1), whereas Troscianko et al. (2017) found up to 11% of variation in human search behaviour to be explained by stimulus appearance when quantified using GabRat. This discrepancy could be explained by the low degree of internal patterning variability close to the edge of each stimulus in our study; a pattern property (edge disruption) that GabRat has been specifically designed to quantify. Thus, despite correlating significantly with detection time, edge contrast metrics in this perceptual context do not appear to capture a single dominating perceptual property of the stimuli driving variation in animal behaviour.

In conclusion, our study highlights the importance of broad and differentiated approaches when concluding ecological relevance from colour pattern statistics. We show the potential of QCPA and its edge detecting statistics to be relevant for the quantification of detection speed and success when considering ecologically relevant viewing contexts. We further provide evidence for a cautious approach towards the identification of pattern statistics responsible for a behavioural response. We acknowledge the many remaining unknowns involved in visual modelling, and we affirm the continued need for ‘context-specific’ behavioural testing of theories and hypotheses brought about by means of theoretical modelling. This, consequently, requires continued testing of and comparisons between colour pattern analyses as they continue to radiate alongside the growing diversity of perceptual and ecological contexts in which they are applied.

## Ethics

All experimental procedures for this study were approved by the University of Queensland Animal Ethics Committee (SBS/077/17).

## Data accessibility

The data are provided here (upload to UQ e-space upon acceptance). All custom Matlab code is available on <u>Github</u>.

## Authors’ contributions

CPvdB: Concept of study, validation, investigation, software, data curation and analysis, original draft, review and editing, project administration, supervision. JAE: Software, validation, review and editing. DEJP: Investigation. KLC: Project funding, validation, resources, review and editing, project administration, supervision.

## Acknowledgements

We would like to thank Nicholas Condon for his assistance with the creation of automated QCPA scripts, various volunteers for assistance with animal husbandry and data entry, as well as the High-Performance Computing (HPC) infrastructure (Wiener & Awoonga) which enabled the computing of image statistics. We would also like to thank two anonymous reviewers for constructive and detailed feedback on the manuscript.

